# Fatty acid synthesis supports tumor progression through keeping TORC1 receptive for Insulin/PI3K signaling

**DOI:** 10.1101/2025.02.19.639068

**Authors:** Dorottya Károlyi, Sólyom Bálint Bótor, Natali Neuhauser, Mária Péter, Gábor Balogh, Fergal O’Farrell, Szabolcs Takáts

## Abstract

Biosynthesis of lipids and fatty acids (FAs) is a tightly regulated and complex process essential for the normal functioning of various cellular processes as is sufficient lipid availability for the progression of several malignant tumor types. Despite its importance, the roles of the individual steps in lipid biosynthesis during tumor growth and their subsequent interaction with intracellular signaling pathways are still not well understood. In our study, we used the Ras^V12^, Scribble deficient carcinoma model in *Drosophila* to demonstrate that upregulation of de novo FA and lipid synthesis is a conserved characteristic of malignant tumors. Performing a small-scale genetic screen by tumor cell specific silencing of components of neutral lipid biosynthetic apparatus revealed that the loss of several enzymes involved in FA and diacylglycerol synthesis significantly inhibited tumor growth. Further characterization of the role of acetyl-CoA carboxylase (ACC), the enzyme responsible for the first step of FA synthesis, revealed that loss of ACC significantly reduced late-stage tumor growth due to increased apoptotic activity. However, early tumor development was unaffected. This correlated with our observation that perturbation of FA synthesis led to inactivation of TORC1 (Target of Rapamycin Complex 1) – the master regulator of cell growth and survival – accompanied by activation of the catabolic process autophagy. Moreover, we also demonstrated that TORC1 activity cannot be restored by hyperactivation of upstream Insulin/PI3K signaling or inhibition of AMP-activated kinase (AMPK) in ACC deficient tumor cells but supplementation of ACC deficient tumors with ectopically added oleic acid alone could improve TORC1 activity and thereby tumor progression. Hence, our findings highlight a new role of FAs in regulating TORC1, rendering it receptive to upstream activatory signals, explaining why cancer cells are extremely dependent on de novo FA synthesis.

## Introduction

Rapidly growing malignant tumors require a large amount of nutrients to maintain the anabolic processes that support the accelerated proliferation of cancer cells. This extreme nutrient demand of tumors eventually leads to the cancer-associated reprograming of the metabolic activity of both the tumor cells and the non-transformed tissues or organs of the host^1,2^. In addition to glucose and amino acids, sufficient lipid availability is also critical for the progression of multiple tumor types. There are two major sources for supporting lipid and fatty acid (FA) demand of tumors. First, lipids can be derived from external sources – from food intake, or can be released by lipolysis from storage tissues, like adipose tissue – and delivered to tumors by circulation. Alternatively, FAs and lipids can be also synthesized de novo from sugars or amino acids by the cancer cells themselves^1^. Although most cells in healthy tissues (except adipose tissues or liver) mainly acquire FAs from external sources instead of synthesizing them, growing body of evidence suggest that cancer cells respond to their increased lipid demand by accelerating their lipid synthetic pathway. Considering that increased FA and lipid synthetic activity is characteristic for multiple tumor types^1,3^, significant efforts have been made to utilize the pharmacological inhibition of lipid biosynthetic pathways as potent anticancer strategy^4,5^.

Lipids represent a large cohort of biochemically diverse biomolecules that have hydrophobic or amphipathic characteristics. Glycerolipids (GLs) are far the most abundant and prevalent lipids that serve as storage lipids (like triacylglycerols/TGs), major components of membranes (glycerophospholipids/GPLs) and are also critical for cellular signaling events (like diacylglycerols/DGs, phosphoinositides) both in normal and cancer cells ^1^. The synthesis of neutral lipids and GPL-s is a complex multistep process, during which the first two carbons of a glycerol-3-phosphate backbone gets esterified with free FAs, while the phosphate group could be modified in various ways: It can be conjugated to organic molecules (like choline, ethanolamine) in various GPLs, it can remain intact (phosphatidic acid/PA), can be removed (like DG) or replaced by another FA (TGs). The biosynthesis of FAs is also a multistep process that begins with Acetyl-CoA that is turned to malonyl-CoA in a reaction catalyzed by Acetyl-CoA Carboxylase (ACC), then this malonyl-CoA is used by the cyclic activity of fatty acid synthase (FASN) to build an elongated acyl chain^1,6^. Although several specific steps of fatty acid (FA) and GL synthesis have been recognized as key processes in tumor physiology, how different enzymes of these synthetic processes and their lipid products regulate cancer cell physiology, especially how cancer cells sense FA and lipid availability and how it influences growth signaling pathways, is still poorly understood.

Insulin/PI3K/TORC1 signaling is the central regulator of lipid synthetic process, which promotes the expression of key enzymes of lipid and FA synthesis like ACC or FASN through the activation of SREBP family transcription factors^7,8^. TORC1 is active upon nutrient rich conditions and promotes anabolic processes (like protein or lipid synthesis) and acts as a critical positive regulator of cell growth^9,10^. Moreover, TORC1 also senses the energetic state of the cells, through AMP-activated kinase (AMPK) that gets activated upon high AMP/ATP ratio and attenuates anabolic processes through inhibitory phosphorylation of TORC1^11^ and ACC^12^. The complex dynamics between ACC activity and Insulin/PI3K/TORC1 signaling is critical to the progression (growth and survival) of cancer cells. Elevated activity of Insulin/PI3K/TORC1 signaling and increased expression of ACC in cancer cells is characteristic for multiple tumor types^1,8,13^. Although genetic or pharmacological inhibition of ACC efficiently attenuates the progression of cultured cancer cells but the cellular and molecular mechanism can vary between different cell types. Loss of ACC in breast and prostate cancer cells results in increased oxidative stress and apoptosis^14,15^, in contrast ACC deficient cholangiocarcinoma cells show decreased proliferation^16^. As the activity of Insulin/PI3K/TORC1 signaling stimulates cell proliferation and inhibits apoptosis^13^, the above-mentioned phenotypes, raises the possibility that loss of FA synthesis may affect cell survival though manipulation of Insulin/PI3K/TORC1 pathway. Importantly, activation of AMPK and inactivation of Akt an upstream member in Insulin/PI3K pathway in ACC deficient cells was reported already^15^, their subsequent effect on TORC1 activity upon these conditions is still poorly understood.

Although experiments on in vitro cultured cells can give good insight how inhibition of FA or lipid synthesis is affecting cancer cell proliferation or death, these setups have limitations recapitulating the real influence of the complex molecular environment provided by host tissues. In recent decades, thanks to its powerful genetic systems that enable tissue specific genetic manipulations in vivo, *Drosophila* became a popular model organism for studying the signaling and metabolic pathways activated in the tumors itself, or in host tissues and how these affect tumor progression in a living organism^17^. It was shown recently that in *Drosophila*, the progression of tumors overexpressing *Ras*^*V12*^ and mutant for cell polarity gene *Scribble* is highly dependent on the import of sugars and amino acids that are released from host tissues by tumor induced catabolic processes^18^. However, whether these tumors also require FA and lipid import to the same extent or they rely more on de novo synthesis is still not understood. Hence in our current research we use the same *Drosophila* tumor model and carry out cancer cell specific inhibition of distinct steps of de novo FA and lipid synthesis to uncover the cell autonomous role of these processes in tumor progression and to reveal how defective FA synthesis is affecting Insulin/PI3K/TORC1 signaling in vivo.

## Results

### 1. Cancer cell specific FA synthesis is required for lipid droplet formation and tumor growth

To characterize the importance of lipid synthesis in the progression of Drosophila tumors we used a well-established fly tumor model and generated malignant tumors in the developing eye (also called eye-antennal disc) of L3 larvae. In this stage the developing eye is composed of rapidly proliferating epithelial cells where we used Mosaic Analysis with a Repressible Cell Marker (MARCM) strategy to generate GFP positive cell clones that overexpress the oncogenic *Ras*^*V12*^ allele and lose their polarity by becoming homozygous mutant for cell polarity gene *Scrib* (hereafter these *Ras*^*V12*^, *Scrib*^*-/-*^ tumors are referred as RS tumors)^17^. First, we aimed to understand whether transformation associated reprograming of lipid metabolism is also characteristic for RS tumor cells. Lipid droplets (LDs) are the most important lipid storage organelles which are composed of neutral lipids and cholesterol esters surrounded by a phospholipid monolayer, and they are present in every healthy imaginal disc cells **(Fig. 1A)**. To understand how the formation and distribution of LDs is changing upon different stages of tumor progression we generated control and FA synthesis deficient RS tumors – by silencing ACC specifically in tumor cells – and stained them with the lipophilic dye monodansylpentane (MDH). Interestingly, we observed that in early stage (day 6 after egg laying) the number of LDs is very low in the GFP positive control tumor tissue but not in GFP negative, non-transformed microenvironmental cells **(Fig. 1B, F)**. Not surprisingly, LDs were completely depleted in ACC deficient cancer cells **(Fig. 1C, F)**, which indicates that transformed cells of early staged tumors may use all the available intracellular storage lipids for their rapid proliferation. In contrast, in mid-late stage (day 7) we observed the reappearance of LDs in multiple control RS tumors **(Fig. 1D, F)**, however LD-s were still lacking in the same aged ACC deficient ones **(Fig. 1E, F)**, indicating that there is an ACC dependent shift in FA and lipid synthesis during transition of tumors from the early to mid-late stages. To assess whether this stage associated dynamics of FA synthesis and LD formation is essential for the growth of this tumor type, we measured the size of control and ACC silenced RS tumors at early (day 6), mid-late (day 7) and late (day 8) stages. Although the loss of ACC decreased tumor size at any stages, but this was only significant in mid-late and late stage **(Fig. 1H-N)**. Hence, ACC mediated FA synthesis and LD formation is required mostly for supporting accelerated progression of RS tumors in later stages, but it is not yet limiting during the growth of early tumor tissues.

**Fig. 1:**
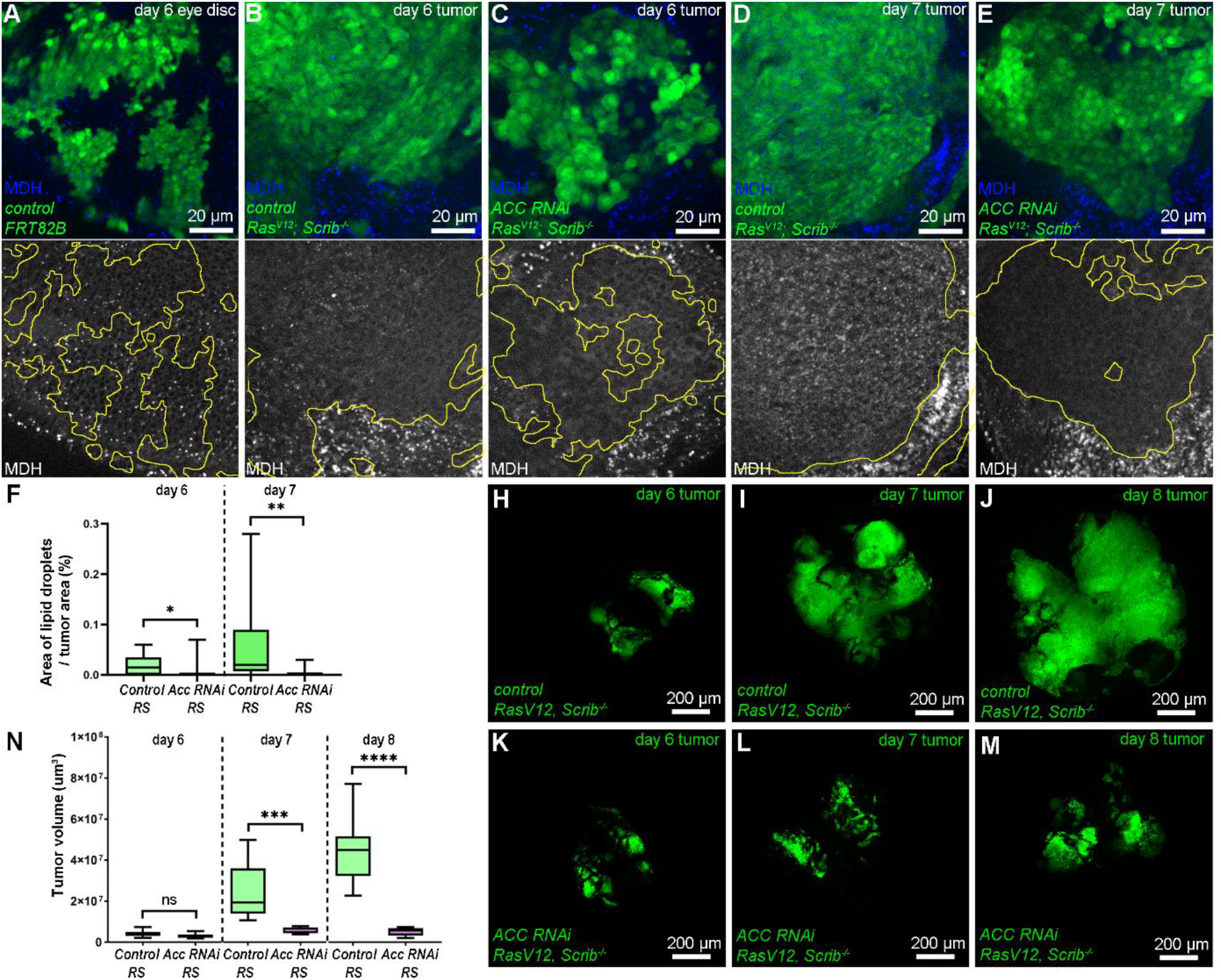
Activation of de novo FA synthesis and LD formation supports the growth of RS tumors. **(A)** LDs are evenly distributed in control eye-antennal discs, and their formation is not affected by generation of control MARCM clones (GFP+). **(B, C)** In early staged (day 6) control RS tumors **(B)** LDs persist in non-transformed microenvironment (GFP-) but mostly disappear from cancer cells (GFP+), and this LD depletion is even more prominent upon cancer cell specific loss of ACC **(C). (D, E)** In mid late stage (day 7), LDs can be detected in control RS tumor cells (GFP+), however their amount is still lower than in the microenvironment (GFP-) **(D)**. In contrast LDs are still undetectable in same staged ACC deficient cancer cells **(E). (F)** Quantification of data presented on B-E. 10-15 tumors/genotypes were analyzed, n=10 (B), 15 (C), 10 (D), 13 (E). Mann-Whitney tests, *: p<0,05; **: p<0,01. **(H-M)** Representative images about early (day 6), mid-late (day 7) and late staged (day 8) control **(H-J)** and ACC deficient **(K-M)** RS tumors. **(N)** Quantification of data presented on H-M. 8-11 tumors/genotypes were analyzed, n=10 (H), 10 (I), 9 (J), 11 (K), 9 (L), 8 (M). Student’s T-test, ns: non-significant; ***: p<0,001; ****: p<0,0001. GFP+ areas are encircled with yellow in grayscale panels of A-E.

### 2. Cancer cell autonomous FA and DG synthesis is critical for tumor progression

Since ACC represents only the very first step of lipid synthesis, while LDs are mostly composed of neutral lipids **(Fig. 2A)**, we carried out a small-scale RNAi screen by knocking down various enzymes required for the production of neutral lipids specifically in the RS cancer cells and assayed their effect on tumor size **(Fig. 2B-E, Supplementary Fig. 1)**. Similarly to ACC, silencing of FASN1 also caused dramatic decrease in tumor size **(Fig. 2C, E)**, which confirmed that RS tumors are highly dependent on de novo FA synthesis. We also silenced several enzymes involved in later stages of lipid synthesis and found that loss of Lpin phosphatidic acid phosphatase, which generates diacylglycerol (DG) from phosphatidic acid (PA), strongly inhibited tumor growth **(Fig. 2D, E)**. In contrast, silencing genes encoding glycerol-3-phosphate acyltransferase (GPAT) enzymes *mino* or *Gpat4* **(Supplementary Fig. 1A, B)**, or the DG acyltransferase (DGAT) *mdy* did not perturb tumor progression **(Supplementary Fig. 1C)**.

**Fig. 2:**
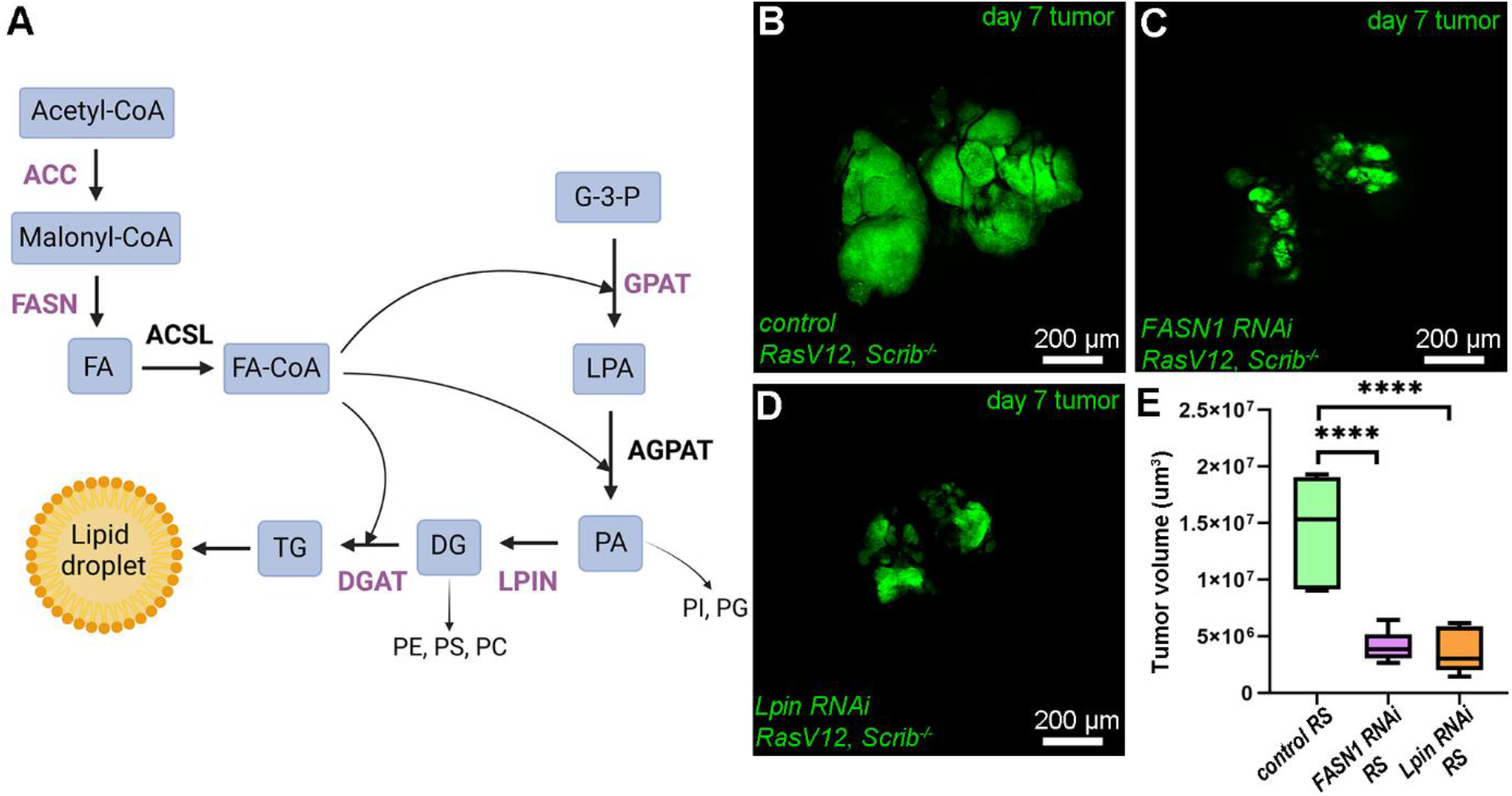
FA and DG synthesis is critical for the progression of RS tumors. **(A)** Schematic figure about the biosynthetic pathway of neutral lipids, by representing lipid classes and organic intermediate products (placed in rectangles) involved in neutral pathway and the major families of lipid biosynthetic enzymes (placed above the arrows). The enzyme families that were tested in the small-scale screen are represented by purple letters. **(B-D)** Representative images at day 7 control **(B)** and FASN1 **(C)** or Lpin **(D)** deficient RS tumors from the RNAi screen. **(E)** Quantification of respective tumor size data represented on B-D. 7-8 tumor/genotypes were analyzed, n=7 (B), 8 (C), 7 (D). Student’s T-test, ****: p<0,0001. FA: Fatty acid; G-3-P Glycerol-3-phosphate; LPA: Lysophosphatidic acid; PA: Phosphatidic acid; DG: Diacylglycerol; TG: Triacylglycerol; PI: Phosphatidylinositol; PG: Phosphatidylglycerol; PE: Phosphatidylethanolamine; PC: Phosphatidylcholine; PS: Phosphatidylserine; ACC: AcetylCoA-Carboxylase; FASN1: Fatty acid synthase 1; ACSL: AcylCoA-Synthase; GPAT: Glycerol-3-phosphate acyltransferase; AGPAT: 1-acylglycerol-3-phosphate-O-acyltransferase; LPIN: Lipin; DGAT: DG acyltransferase.

### 3. *de novo* FA synthesis protects tumor cells from apoptosis

Rapid proliferation and the appearance of large apoptotic areas – that are positive for active caspases – laying mostly in the microenvironment is characteristic for RS tumors^19^. However, the impaired progression of RS tumors deficient for FA synthesis brought up the question whether these tumors cannot grow due to a proliferation defect or upregulation of cell death. To assess the intensity of these two processes we carried out immunolabelling on day 6 tumors where the area of the transformed and microenvironmental tissues are comparable. By staining the nuclei of proliferating cells by Phospho-Histone H3 (P-H3) we did not observe any difference between cancer cell clones of control or ACC deficient RS tumors in the frequency of mitotically active cells **(Fig. 3A-C)**. In contrast, assaying apoptotic areas by carrying out immunolabelling of Cleaved Death Caspase-1 (Dcp-1) revealed that this apoptotic marker overlaps with ACC deficient cancer cell clones to a significantly larger extent than it does in controls **(Fig. 3C-D)**. Hence, elevated FA synthesis may act as a cytoprotective process that helps cancer cells to evade apoptosis.

**Fig. 3:**
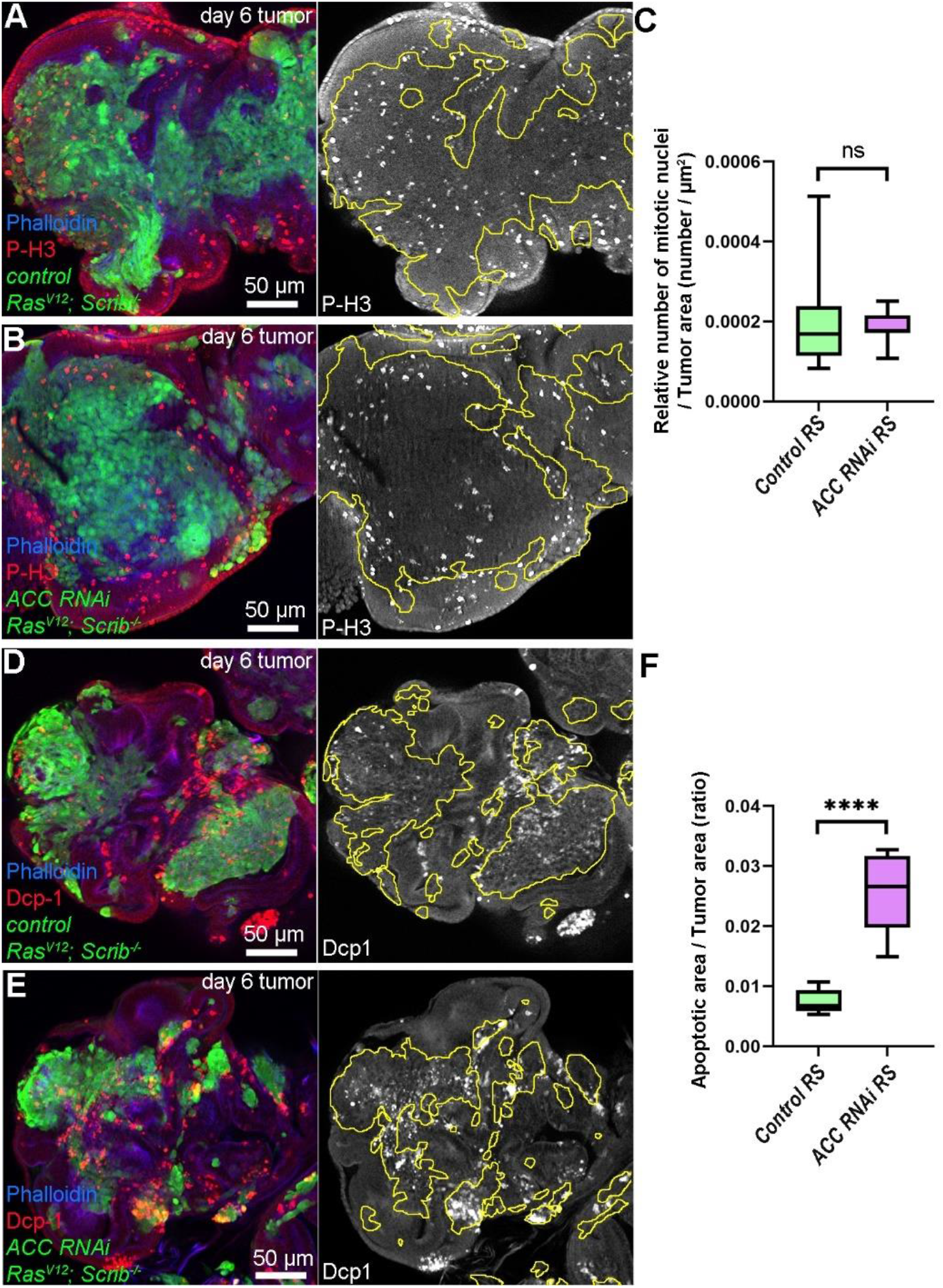
Loss of ACC promotes cancer cells’ death but does not affect their proliferation. **(A, B)** Comparison of control **(A)** and ACC deficient **(B)** tumor (GFP+) in the relative number of mitotically active cells (positive for anti-P-H3 staining) compared with microenvironmental cells (GFP-). **(C)** Quantification of data presented on A, B. 9-10 tumors/genotypes were analyzed, n=10 (A), 9 (B). Mann-Whitney test, ns: non-significant. **(D, E)** Immunolabeling of active effector caspase Dcp-1 for comparison of ACC deficient tumor tissue (GFP+) or control to microenvironment (GFP-). **(F)** Quantification of data presented on D, E. 6 tumors/genotypes were analyzed, n=6 (D, E). Student’s T-test, ****: p<0,0001. GFP+ tumor areas are encircled with yellow in grayscale panels of A, B, D, E.

### 4. Tumor specific loss of ACC reduces TORC1 activity and makes it poorly inducible by Insulin/PI3K pathway

Insulin/PI3K/TORC1 pathway induces cell growth, proliferation and lipid synthesis, however it is also known as a negative regulator of apoptosis ^13^. Considering the weak growth potential and elevated apoptotic rate of ACC deficient RS tumors we wondered whether loss of FA synthesis may perturb TORC1. Active TORC1 not only promotes translation by phosphorylating S6-Kinase (S6K) and elongation initiation factor 4E binding protein (4E-BP), but also inhibits the lysosomal catabolic process, autophagy. Carrying out immunostainings with antibodies specific for P-S6K revealed that cancer cells in both early and mid-late staged control RS tumors show elevated TORC1 activity that is indicated by increased P-S6K signal compared to the microenvironment **(Fig. 4A, C)**. In contrast, the upregulation of P-S6K was not apparent in ACC deficient cancer cells **(Fig. 4B, D)**, indicating the activity of TORC1 is reduced in these tumors. Assaying autophagic activity by immunolabelling autophagosome marker Atg8a led to the same conclusion. While Atg8a signal was mostly cytoplasmic and only a few autophagic structures appeared in control tumors **(Fig. 4E, G)**, ACC deficient cancer cells showed the accumulation of Atg8a positive autophagosomes **(Fig. 4F, G)**. As further confirmation that cancer cell specific loss of ACC impairs TORC1 activity we carried out western blots on the lysates of late staged tumors and we observed ACC silenced tumors show massive decrease in the levels of phosphorylated form of 4E-BP **(Fig. 4H)**.

**Fig. 4:**
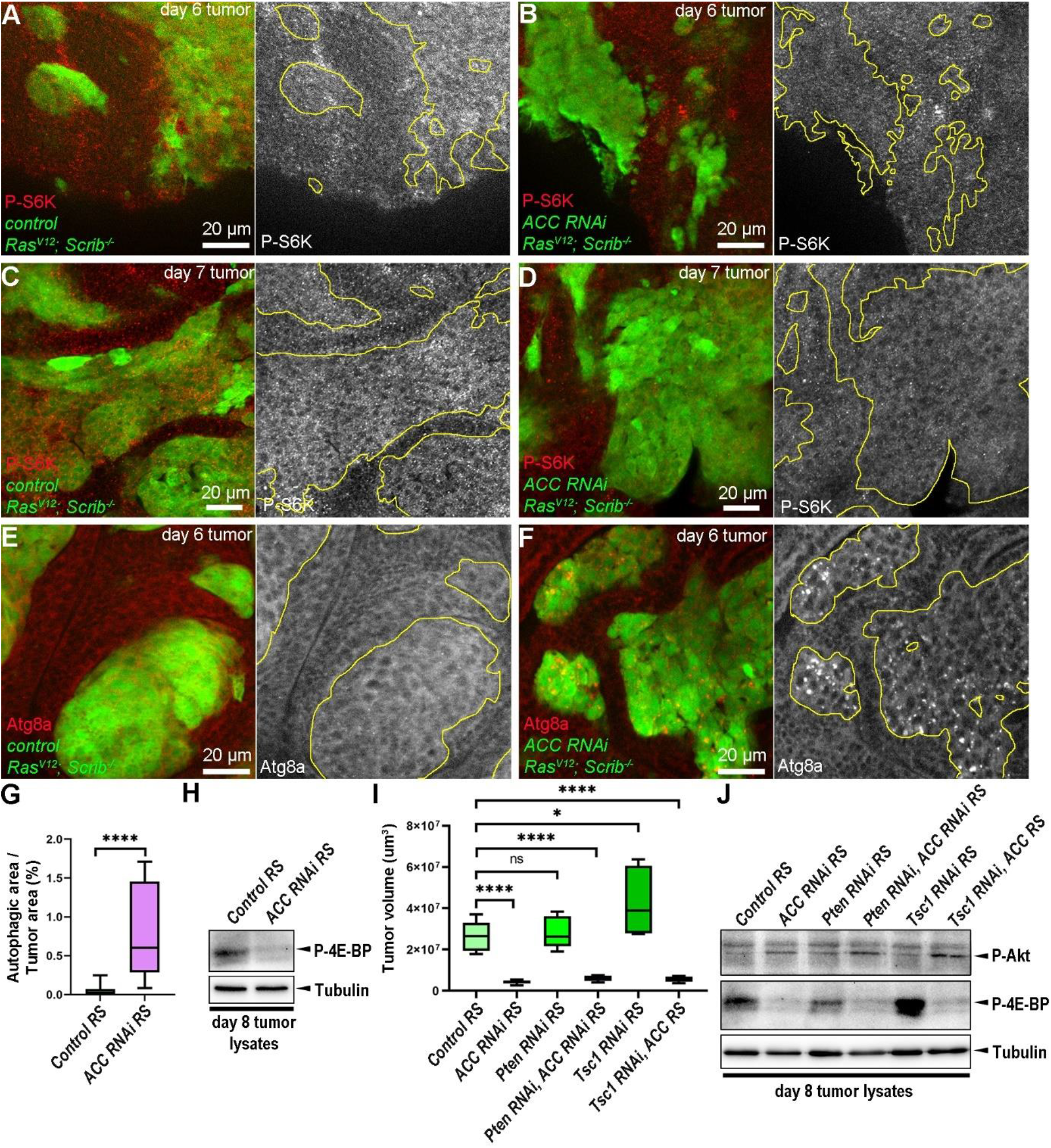
Tumor cell specific loss of ACC inactivates TORC1 and makes it non-receptive for incoming signals of Insulin/PI3K pathway. **(A-D)** Anti-P-S6K immunolabelling to investigate TORC1 activity in early (day 6) and mid-late staged (day 7) control **(A, C)** and ACC deficient **(B, D)** tumors (GFP+) and their microenvironment (GFP-). **(E, F)** Immunolabelling with anti-Atg8a reveals the number and distribution of autophagic structures in control **(E)** and ACC RNAi **(F)** RS tumor cell clones (GFP+). **(G)** Quantification of data presented on E, F. 10-13 tumors/genotypes were analyzed, n=13 (E), 10 (F). Mann-Whitney test, ****: p<0,0001. **(H)** Anti-P4E-BP western blot on lysates of late staged (day 8) control and ACC deficient RS tumors. **(I)** Tumor size analysis on day 8 tumors reveals how cancer cell specific loss of Pten and Tsc1 is affecting the growth potential of control and ACC deficient RS tumors. 6-10 tumors/genotypes were analyzed, n=8 (control RS), 10 (ACC RNAi RS), 8 (Pten RNAi RS), 6 (Pten RNAi, ACC RNAi RS), 7 (Tsc1 RNAi RS), 9 (Tsc1 RNAi, ACC RNAi RS). Student’s T-test, ns: non-significant; *: p<0,05; ****: p<0,0001. **(J)** Western blotting on day 8 tumors to reveal the effect of loss of Pten or Tsc1 on the levels of P-4E-BP and P-Akt in control and ACC deficient tumors. Tubulin represents loading control in H and J. GFP+ tumor areas are encircled with yellow in grayscale panels of A-F.

As decreased TORC1 activity seemed to be a likely cause of diminished progression of ACC deficient tumors, we supposed that the growth of these tumors might be partially restored by reactivating TORC1 through ectopic induction of Insulin/PI3K pathway. Hence, we generated tumors that simultaneously silenced ACC and known negative regulators of Insulin/PI3K pathway, Phosphatase and tensin homolog (Pten) and Tuberous sclerosis 1 (Tsc1). Reducing levels of Pten and Tsc1 caused a mild overgrowth in both RS and RS ACC conditions, although comparisons between RS and RS ACC show no significant rescue of the tumor growth phenotype, remaining tiny in comparison to RS **(Fig. 4I)**. Western blot experiments confirmed that silencing of Pten or Tsc1 in ACC deficient tumors had only a minimal if any positive effect on TORC1 activity (P-4E-BP levels), indicating that activation of Insulin/PI3K signaling was inefficient in stimulating TORC1 in these samples **(Fig. 4J)**. This result suggests that the low growth potential of ACC deficient RS tumors is due to constantly blunted TORC1 activity. This is further highlighted when we compare these results with control RS tumors where silencing of Tsc1 caused massive induction of TORC1 activity that is indicated by significant increase in tumor size and striking accumulation of P-4E-BP **(Fig. 4I, J)**. Interestingly, the levels of P-Akt – the active form Akt, a member of Insulin/PI3K/TORC1 pathway upstream from Tsc1 – was higher in ACC silenced tumors even in the presence or absence of PTEN or Tsc1 **(Fig. 4J)**. This suggests that in contrast to TORC1, the more upstream steps of Insulin/PI3K pathway are already upregulated upon the loss FA synthesis, likely explaining why ectopic induction of Insulin/PI3K pathway have a minimal stimulatory effect on TORC1 activity in these tumors. This highlights that FA synthesis may strictly regulate TORC1 by making it responsive for upstream stimulatory signals from Insulin/PI3K pathway.

### 5. Upregulation of AMPK plays a minor role in TORC1 inactivation in ACC deficient tumors

In addition to Insulin/PI3K, TORC1 can be stimulated by the intracellular levels of nutrients and ATP, and low levels of these are activating AMPK that stalls anabolic processes through direct inhibition of TORC1 by phosphorylation, or indirectly through activation of Tsc1. To test whether AMPK is activated upon the loss of ACC we immunolabelled the active phosphorylated form of AMPK in RS tumors. Cancer cells of control RS tumors showed suppressed P-AMPK signal compared to the non-transformed microenvironmental cells **(Fig. 5A)**. In contrast ACC deficient cancer cells did not show such a decrease in P-AMPK levels **(Fig. 5B)**. This finding suggested that AMPK is upregulated in cancer cells with impaired FA synthesis. To test whether this effect can be responsible for the inactivation of TORC1 in ACC silenced tumors we generated RS tumors double deficient for ACC and the α-subunit of AMPK. However, and somewhat surprisingly, western blotting revealed that inactivation of AMPK had only a minimal stimulatory effect on P-4EP-BP levels **(Fig. 5C)**, and did not restore the size of ACC deficient tumors in comparison to control RS **(Fig. 5D)**.

**Fig. 5:**
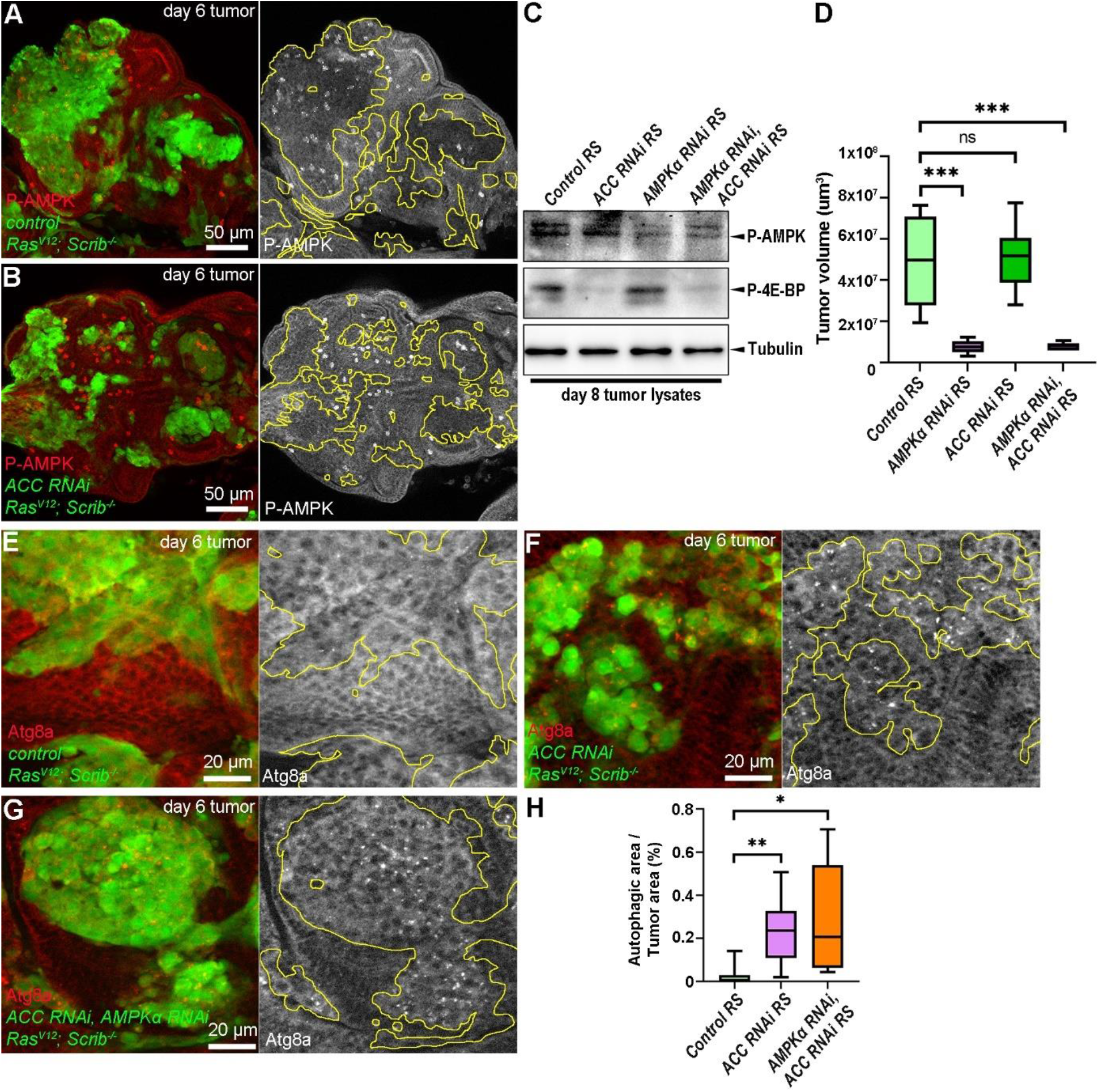
Activation of AMPK is not responsible for diminished progression of ACC deficient RS tumors. **(A, B)** Anti-P-AMPK immunolabeling in control RS **(A)** and ACC deficient **(B)** tumor cells (GFP+) and in their microenvironment (GFP-). **(C)** Western blotting of day 8 tumor samples indicates ACC and AMPKα RNAi mediated changes in the levels of P-4E-BP and P-AMPK. **(D)** Analysis of the size of day 8 tumors to assay the influence of the presence and the loss of AMPK on control RS and ACC deficient tumors. 6-10 tumors/genotypes were analyzed, n=7 (control RS), 8 (ACC RNAi RS), 10 (AMPKα RNAi RS), 6 (AMPKα RNAi, ACC RNAi RS). Student’s T-test, ns: non-significant, ***: p<0,001. **(E-G)** Immunolabeling reveals that autophagic structures – that are missing from control **(E)** but accumulating in ACC deficient **(F)** RS tumor cells (GFP+) – are still present in AMPKα, ACC double deficient **(G)** tumor tissues (GFP+). **(H)** Quantification of data presented on E-G. 5-9 tumors/genotypes were analyzed, n=7 (E), 9 (F), 5 (G). Mann-Whitney test, *: p<0,05; **: p<0,01. GFP+ tumor areas are encircled with yellow in grayscale panels of A, B, E-G.

As AMPK can also induce catabolism by stimulating autophagy through inactivation of TORC1 or direct phosphorylation of Atg1/ULK1 complex^11^ we questioned whether activated AMPK is responsible for the above-described autophagy induction in ACC silenced RS tumor cells **(Fig. 4F, G)**. However, immunolabelling of autophagic structures revealed that concomitant inhibition of AMPK is not decreasing the amount of Atg8a positive autophagosomes in ACC deficient cells **(Fig. 5E-H**), suggesting that the induction of autophagy in these tumors is independent of AMPK. Taking these together our findings imply that although ACC deficient cancer cells show elevated AMPK activity, it is likely that TORC1 inhibition in these tumors is driven by AMPK independent means.

### 6. Fatty acids can activate TORC1 in ACC deficient RS tumors

Since neither the activation of Insulin/PI3K pathway nor inhibition of AMPK could stimulate TORC1 in ACC deficient RS tumors, we supposed that the availability of FAs or some lipid species may have a direct role in the maintenance of TORC1 activity in cancer cells. To assay how the lipid composition is changed upon the loss of FA or lipid synthesis we carried out shotgun lipidomics analysis on late staged (day 8) control and ACC and Lpin deficient RS tumors **(Supplementary Table S1)**. This way we could distinguish between the common and specific alterations in the lipid profile that were caused by cancer cell specific perturbation of FA (*ACC RNAi*) or DG (*Lpin RNAi*) synthesis. Most interestingly, comparing the lipid profiles of the three genotypes by Cluster **(Fig. 6A)** and Principal component analysis **(Fig. 6B)** revealed that although both ACC and Lpin deficient samples sharply diverge from controls or from each other, Lpin RNAi samples still positioned somewhat closer to controls than ACC RNAi, pointing out that loss of FA synthesis causes a more substantial alteration in the lipid composition of cancer cells. Importantly, loss of ACC caused a general alteration in the fatty acid profiles with a significant decrease in lipid species composed of mostly saturated or monounsaturated FAs, while lipids that contained at least one polyunsaturated FAs (PUFAs, with two or more double bonds) were accumulated **(Fig. 6A, C, D; Supplementary Fig. S2)**. Considering that the enzymatic apparatus for the synthesis of PUFAs is lacking from *Drosophila* genome, these PUFAs can only be derived from food, while saturated and monounsaturated lipids can be synthetized de novo. Hence, this FA composition reveals a massive decrease in de novo synthetized FAs in ACC deficient RS tumors.

**Fig. 6:**
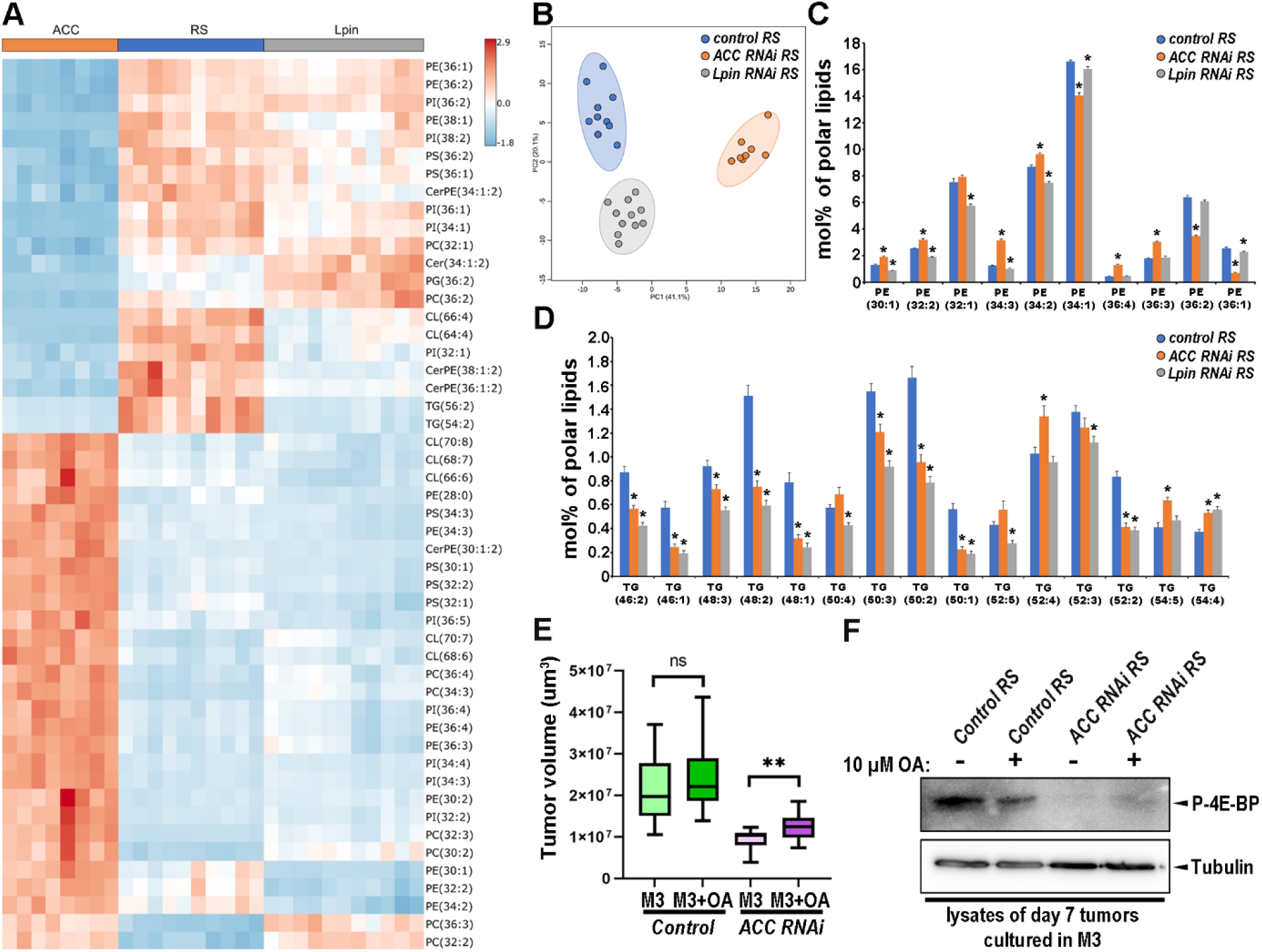
Saturated and monounsaturated FAs are critical for the progression of ACC deficient tumors. **(A-D)** Shotgun lipidomics analysis of day 8 control, and ACC or Lpin deficient RS tumors. Relative amounts were measured and normalized for polar lipids. **(A)** Cluster analysis representing top 50 lipid species whose amount produces the highest relative changes in the analysis. **(B)** Principal Component Analysis reveals that lipid profiles of ACC RNAi samples are more divergent from controls than Lpin deficient ones. **(C, D)** Diagrams representing the relative amount of the most important PE (those that are above 1%) and TG species (those that are above 0,2%). Analysis of PE profile indicates that species composed of mostly long chained saturated or monounsaturated FAs (like PE 34:1; PE 36:1 and PE 36:2) are decreased, while those ones with 3 or 4 double bonds (that are containing at least one polyunsaturated FA) are significantly upregulated in ACC deficient tumors **(C)**. Analysis of TG profile shows that most of the TG species are strongly decreased in both ACC and Lpin RNAi tumors. However, those TG-s that contain at least one polyunsaturated FA (4 or more double bonds) are also significantly upregulated in ACC deficient samples **(D)**. Statistical analysis on C and D was done by comparing values from ACC RNAi RS and Lpin RNAi RS to control RS data by using Student’s T-test and asterisk (*) above the columns represent significant difference (p<0,05). 8-11 samples/genotypes were analyzed, n=10 (control RS), 8 (ACC RNAi RS),11 (Lpin RNAi RS). (**E)** Analysis of the size of day 7 tumors that were previously cultured in ex vivo conditions in normal M3 or oleic acid (OS) supplemented M3 media for 24 h. 10-19 samples/genotypes were analyzed, n=10 (control RS M3), 10 (control RS M3+OA), 18 (ACC RNAi RS M3), 19 (ACC RNAi RS M3+OA). Student’s T-test (control RS M3 vs control RS M3+OA) and Mann-Whitney test (ACC RNAi RS M vs ACC RNAi RS M3+OA), ns: non-significant, **: p<0,01. **(F)** Western blotting to measure the levels of P-4E-BP in the lysates of day 7, ex vivo cultured tumors treated with or without OA.

In contrast to ACC RNAi tumors the genotype specific changes of Lpin deficient samples are affecting only some specific lipid species, like significantly increasing PG and decreasing PE levels **(Fig. 6A, C, Supplementary Fig. S2A, B, F)**. Finally, we observed several changes in lipid profile that were common in both ACC and Lipin deficient samples compared to controls. One such common feature was the significant decrease in the relative amount of TG species that contain saturated or monounsaturated FAs (like TG(48:2)) **(Fig. 6D, Supplementary Fig. S2A, H)**, which indicates a general decrease in de novo lipid synthesis in both genotypes and it is in line with our observation about the lack of LDs from ACC deficient cancer cell clones. Additionally, we also observed significant decrease in the relative amount of phosphatidylserine (PS) and ceramide-phosphoethanolamine (CerPE), and a remarkable increase in phosphatidylcholine (PC) both in ACC and Lpin deficient tumors **(Fig. 6A, Supplementary Fig. S2A, C)**, which might represent general characteristics of cancer cell specific rearrangement of lipid synthesis.

Considering that the deficit in saturated and monounsaturated FAs is the most characteristic alteration in the lipid profile of ACC silenced tumors, we aimed to examine whether this feature can be a key for inactivation of TORC1 in these tumors. Hence, we designed an experimental setup where we maintained tumors dissected on day 6 larvae in Insect M3 media for 24 h. This setup enabled us to directly supplement ex vivo cultured tumors with fatty acids and monitor effects to tumor progression. ACC deficient RS tumors that were incubated in M3 media supplemented by 10 μM oleic acid (OA) grew significantly larger than those that were incubated in pure M3 **(Fig. 6E)**. Importantly, this effect of OA supplementation did not affect the growth potential of ex vivo cultured control RS tumors **(Fig. 6E)** (which presumably were capable of de novo synthesis). Finally, western blotting also revealed that treating ex vivo cultured ACC silenced RS tumors with M3 + OA but not M3 alone increases P-4E-BP levels, an effect again not observed in FA de novo competent control RS tumors **(Fig. 6F)**. These findings shed light on a mechanism where de novo synthetized FAs directly regulate TORC1 activity, likely by keeping TORC1 receptive for incoming upstream stimulatory signals from Insulin/PI3K pathway.

## Discussion

In our study we demonstrated that like multiple human tumors the *Drosophila* RS tumors are also undergoing a metabolic shift that includes accelerated de novo synthesis of FAs and lipids crucial for tumor progression. We found that cancer cell specific knock down of ACC, FASN1 or Lpin enzymes, diminished tumor growth and we provided evidence that induction of lipid synthesis in cancer cells is vital for their survival and progression through TORC1 regulation. Importantly, these findings also shed light on the high demand on FAs and lipids that these tumors experience, which cannot be fulfilled by food intake or the lipolytic capacity of the host microenvironmental tissues. Although not all of the lipid synthetic enzymes were found to be essential for tumor growth in our genetic screen, this discrepancy can be likely resolved by the fact that these enzymes have multiple paralogs in *Drosophila* genome (three GPAT and four DGAT family genes are listed in FlyBase) and may act redundantly, while Lpin represents the only PA phosphatase ortholog. Hence, our results point out the importance of cancer cell specific de novo FA and DG synthesis in supporting RS tumor formation, however we cannot exclude that GPAT or DGAT functions may also contribute to this process. The importance of de novo FAs for tumor progression was also demonstrated by shotgun lipidomics that indicated a massive decrease of de novo synthetized FAs among the components of membrane lipids and TGs in ACC but not Lpin deficient tumors. However, it is important to note that the larvae we investigated were raised on normal diet, and we cannot exclude that rearing larvae on lipid-rich media may partially mitigate the deleterious effect of the lack of ACC on tumor growth. This scenario is further strengthened by our ex vivo experiment where we observed that the progression of ACC deficient tumors could be improved by incubating them in OA supplemented media.

Our findings confirmed that tumor specific downregulation of ACC is decreasing tumor size through increasing cancer cell apoptosis instead of decreasing their proliferation. These findings are in line with studies about human cancer cell lines, derived from head and neck squamous carcinoma, prostate cancer and breast cancer^14,15,20^. Although some other studies also report decreased proliferation of cancer cell lines that are deficient for ACC^16^, we did not observe any such effect in *Drosophila* tumors at the age we tested (day 6). However, we cannot rule out the possibility that silencing of ACC may affect cancer cell proliferation in later staged RS tumors.

Our observation that TORC1 activity is diminished in ACC deficient tumors offers a clear-cut explanation why these cells show increased apoptotic activity. Moreover, we also demonstrated that loss of ACC results in increased activity of AMPK and induces autophagy. Although other studies also reported that ACC inhibition is accompanied by elevated AMPK activity and lower Insulin/PI3K signaling^15,16^, those studies have not really tested whether these changes are causes or just byproducts of decreased progression of ACC deficient cells. One of the advantages of our *Drosophila* model was that it enabled us to carry out rescue experiments. In this manner we could demonstrate that either reactivation of Insulin/PI3K signaling, or silencing of AMPK, only had minimal effects in restoring TORC1 activity and growth of ACC deficient tumors. This indicates that loss of ACC makes TORC1 insensitive for signals from upstream members of Insulin/PI3K pathway and AMPK. Our findings are in line with a previous *Drosophila* paper, which showed that mutation of FASN1 can decrease the overgrowth of Pten deficient but not normal adipose tissue cells^21^. This suggests that accelerated FA synthesis may be required for sustaining TORC1 responsiveness for Insulin/PI3K signaling when the pathway is hyperactivated. Considering that oncogenic Ras^V12^ can also activate PI3K^17^, it is likely that Insulin/PI3K pathway is also hyperactivated in most tumors with elevated Ras activity (like the RS tumors) and de novo FA synthesis may sustain TORC1 to be recipient for these stimulatory signals **(Fig. 7)**, hence inhibiting FA synthesis might be a particularly efficient anti-tumor therapeutic strategy against Ras tumors. Moreover, this strategy has further a benefit as it may affect TORC1 activity mostly in cancer cells rather than in the healthy ones.

**Fig. 7.**
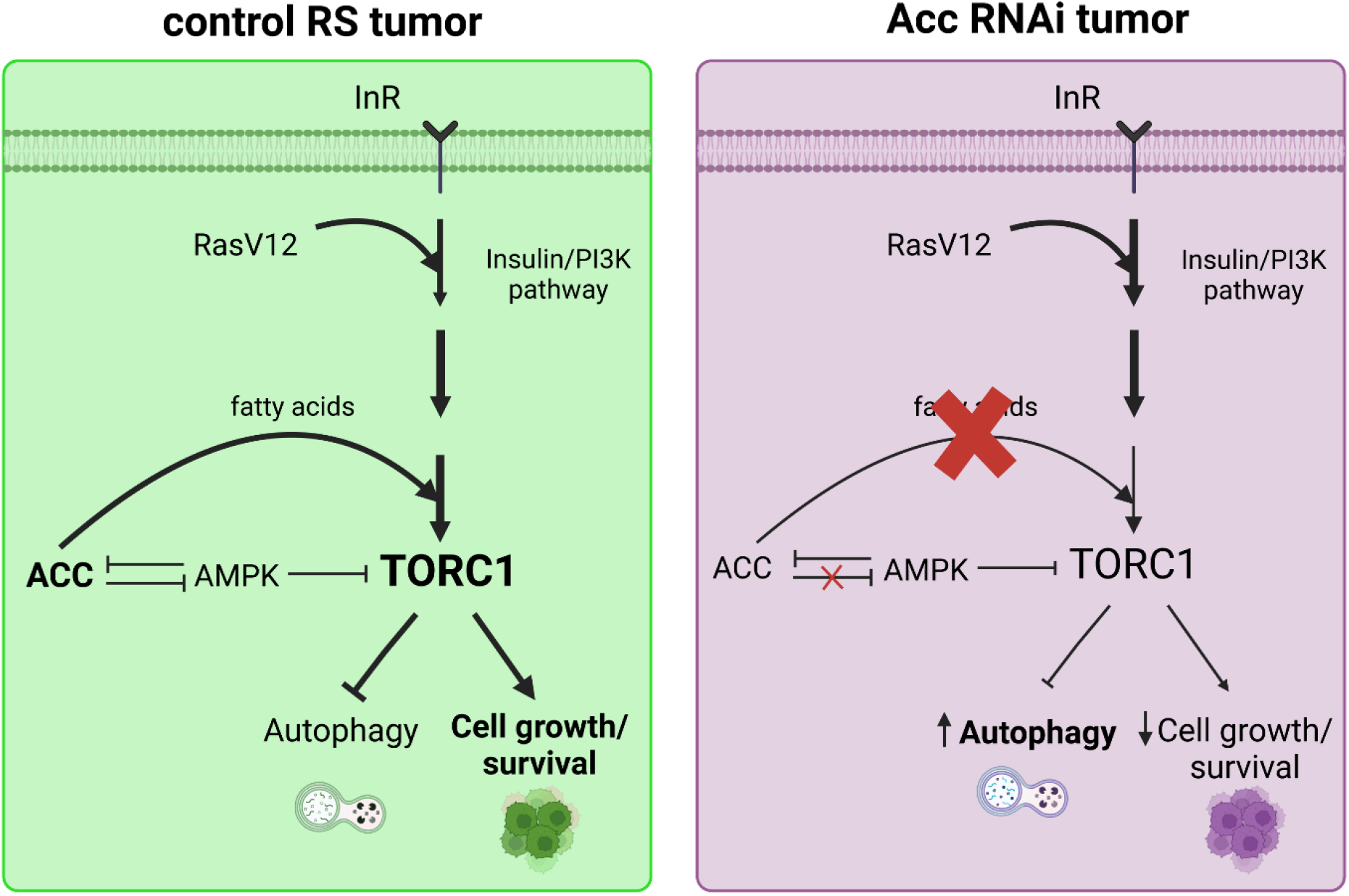
FA synthesis promotes TORC1 activity by keeping it receptive to Insulin/PI3K pathway derived signals in cancer cells. In our proposed model RS tumors can rapidly progress due to at least two effects of Ras^V12^. First Ras^V12^ directly induces proliferation through MAPK pathway. Second Ras^V12^ can also stimulate TORC1 mediated cell growth and evasion of apoptosis through ectopic activation of Insulin/PI3K pathway. Our findings pointed out that high levels of TORC1 activity could be maintained in the presence of ACC mediated de novo FA synthesis, which keeps TORC1 receptive to the incoming upstream signals from the Insulin/PI3K pathway – through a yet unknown mechanism –, and to a minor extent by moderating AMPK activity. In contrast, the lack of ACC results in critical remodeling in the FA composition of cellular lipids, which eventually inactivates TORC1 and stimulates autophagy. Since Ras^V12^ and activated Insulin/PI3K pathway is still present in these cells, the depletion of TORC1’s activity in ACC deficient cancer cells cannot be explained by simply the loss of upstream stimulatory signals. Instead, this situation can be rather interpreted such that TORC1 cannot respond to the upstream stimulatory signals upon the lack of FAs. Hence, our model suggests that de novo lipid synthesis has a critical role in tumor progression by facilitating TORC1 to be receptive for elevated Insulin/PI3K signaling.

The molecular mechanism by which FAs facilitate the responsiveness of TORC1 for upstream factors of Insulin/PI3K pathway is still puzzling. Our ex vivo experiments showing that addition of OA to ex vivo cultured ACC deficient tumors alone was enough to increase TORC1 activity and improve their progression further suggest that FAs may have a direct effect in promoting TORC1 activity. In addition to Insulin/PI3K pathway, TORC1 activity can be also regulated by its subcellular localization, and it was shown previously that palmitate treatment can activate TORC1 in cultured podocytes through inducing the relocalization of TORC1 to the lysosomal membranes^22^. Although it is a tempting possibility that loss of ACC may downregulate TORC1 activity through disrupting its subcellular localization in tumor cells, this hypothesis needs experimental evaluation in a system where TORC1 localization can be tracked efficiently.

Based on our findings, increased autophagy is characteristic for cancer cells lacking ACC. Rescue experiments also elucidated that ACC, AMPK double deficient tumor cells still showed high autophagic activity, indicating that induction of autophagy in these cells is likely due to TORC1 downregulation rather than to the elevation of AMPK activity. We can assume that autophagy may have a protective function in cells defective for FA synthesis because autophagy can release FAs by the degradation of dispensable or damaged membrane bound organelles.

This raises the possibility that autophagy may have a critical function in keeping the remaining potential for survival and proliferation of ACC deficient cells and inhibition of autophagy may further improve the efficiency of small molecule inhibitors of ACC, which are already considered as promising anti-tumor drugs^4,5^. However, elucidating whether autophagy has this pro-survival function in ACC deficient cancer cells remains to be determined.

In summary, our study highlights that efficient de novo FA synthesis is required in order to keep TORC1 responsive to the altered incoming growth signals experienced in tumor cells. Furthermore, inhibition of FA synthesis might be an efficient anti-tumor strategy in tumors characterized with elevated activity of Ras and Insulin/PI3K signaling pathways.

## Materials and Methods

### Drosophila genetics

During the experiments, fruit flies were maintained in glass tubes on agar-corn flour-yeast medium, at constant 25°C.

Malignant tumors were induced using the MARCM (Mosaic Analysis with Repressible Cell Marker) strategy. by using an *ey-Flp, UAS-Dcr; Act-FRT-CD2-FRT-Gal4, UAS-GFP; FRT82B tub-Gal80* genotype that were crossed with various tumor effector lines that carried a transgenic *UAS-Ras*^*V12*^ oncogene, and loss of function mutant allele of *Scribble* (*Scrib*) cell polarity gene (*Scrib*^*1*^).

All mentioned transgenic and mutant stocks were obtained from Bloomington Drosophila Stock Centre (Indiana, US) except *UAS-ACC[KK102082]* and *UAS-FASN1[GD29349]* RNAi lines that were ordered from Vienna Drosophila Research Centre. The detailed description of the used genotypes regarding each experiment are listed in **Supplementary Table S2**.

### Lipid staining

From all genotypes, 6-8 eye discs/tumors of 6 and 7-day-old larvae were dissected in PBS solution and fixed in 4% formalin-PBS for 30 minutes on rotary shaker. To remove the fixative, samples were rinsed twice and washed for 15 minutes in PBS buffer. Then samples were incubated in PBS containing monodansylpentane/AUTODOT (Abcepta) fluorescent lipophilic dye (with 1:500 dilution) overnight at 4°C on orbital shaker. The next day, the staining solution was removed by 2x rinsing and 15 minutes washing in PBS. Finally, the samples were mounted in glycerol and imaged by Zeiss AxioImager M2 fluorescence microscope equipped by ApoTome2 (Zeiss) confocal unit and Orca-Flash 4.0 LT3 digital sCMOS camera (Hamamatsu Photonics).

### Tumor volume analysis

Larvae were raised under standardized conditions to ensure consistency across samples. All genetic crosses were maintained in the same tray, flipped at the same time, and dissected at a consistent time of day and control tumors were always collected in parallel to the RNAi treated ones. This way each experiment had an internal control which mitigates the environmental variance. Tumors were dissected from larvae with the appropriate age in PBS and fixed in 4% formalin-PBS for 30 minutes. To remove the fixative, samples were rinsed two times and washed for 15 minutes in PBS. Then tumors were mounted in glycerol. To avoid the compression of the tumors, spacers (with thickness of 0,2 mm) were placed between the glass slide and the coverslip. The whole volume of tumors was scanned through by optical sectioning (10x magnification Z-stack function, 3-micron slice intervals) by using the already mentioned Zeiss AxioImager M2 fluorescence microscope.

### Immunolabellings

On the first day of fluorescent immunolabelings, 6-8 developing eyes were dissected from 6-or 7-day-old larvae of RC, RS and RS ACC genotypes and fixed in 4% formalin-PBS solution for 20 minutes. After washing three times for 5 minutes each in 0.5% Triton X100-PBS solution (PBT) in order to remove the fixative, primary antibodies **(Supplementary Table S3)** were added in PBT containing 5% fetal bovine serum (FBS PBT). The samples were then incubated overnight at 4°C on orbital shaker. On the second day, after washing three times for 5 minutes in PBT, samples were incubated with secondary antibodies dissolved in FBS PBT for 3 hours at room temperature, in dark chamber. After removing the secondary antibodies, samples were washed two times in PBT for 5 minutes and then twice in PBS for 5 minutes. The samples were then mounted in glycerol and a Z-stack was acquired for each sample using a fluorescence microscope. For P-H3, Dcp1, and p-AMPK immunostainings, images were taken with 1.35 μm (optimal) slice intervals at 20x magnification, while for p-S6K and Atg8 immunostainings, images were taken with 0.35 μm (optimal) slice intervals at 40x magnification.

### Ex vivo culturing

The following incubation media was prepared and warmed to room temperature before the experiment: Shields and Sang’s M3 insect medium (Merck) that contained Penicillin-Streptomycin antibiotics (diluted in 1:100) and optionally supplemented by 10 μM oleic acid (#O3008, Merck). To prevent any contamination sterile 24-well plate was used for all experiments and larvae and forceps tips were washed in 70% ethanol before dissection. For the assay, control RS and ACC RNAi RS tumors were dissected in M3 medium, and 10-15 tumors per genotype were incubated in 500-500 μl of normal or oleic acid containing M3 medium for 24 hours at 25°C, protected from direct light. After incubation, tumor sizes were examined using the above-described volumetric measurement protocol.

### Western blot

Tumors from 8 days old larvae were collected in Eppendorf tubes containing lysis buffer (50 mM Tris-HCl, 150 mM NaCl, 1% Triton X-100, and 5 mM EDTA) supplemented with protease (PMSF 1:100, leupeptin 1:1000, aprotinin 1:1000) and phosphatase (NaF 1:100, Na^3^VO^4^ 1:100) inhibitors and were kept on ice. The samples were homogenized with plastic pestles and their protein concentration was measured by standard Bradford reaction. Finally, after the addition of Laemmlii loading buffer, the samples were boiled for 5 minutes and stored at -20^o^C until further use.

In western blot analysis, samples with equal amounts (15 μg/sample) of protein were loaded to 13% polyacrylamide gel, and after the electrophoresis (Mini-Protean Tetra Cell device, Bio-Rad) the separated proteins were transferred to a PVDF membrane by using a Trans-Blot Turbo semidry blotting device (Bio-Rad). The membrane was washed three times in TBS buffer containing 0.1% Tween-20 detergent (TBST) for 5 minutes each, followed by a 1-hour blocking step in 0.5% casein. Then, the membrane was incubated overnight in a 1:1 TBST-Casein solution containing different primary antibodies **(Supplementary Table S3)** at 4°C. Next day, the primary antibody was removed by 3x washing for 10 minutes in TBST. Then, the membranes were incubated for one hour at room temperature in 1:1 TBST-Casein that contained horseradish peroxidase (HRP)-conjugated secondary antibodies. Secondary antibodies were removed by 3×10 minute wash with TBST and membrane was developed by chemiluminescent HRP substrate solution and imaged by using a Chemidoc Imaging System (Bio-Rad) device.

### Shotgun lipidomics

To minimize contamination, day 8 larvae were first rinsed in ethanol, and then washed in PBS. Three tumors/sample were dissected in sterile PBS and were carefully collected using dissection needles to minimize excess fluid and immediately transferred into empty Eppendorf tubes, which were then flash-frozen in liquid nitrogen. Then samples were kept on -80°C until further analysis.

Lipidomic standards were from Avanti Polar Lipids (Alabaster, AL, USA). Solvents for extraction and mass spectrometry (MS) analyses were Optima LC-MS grade (Thermo Fisher Scientific, Waltham, MA, USA) and liquid chromatographic grade (Merck, Darmstadt, Germany). All other chemicals were the best available grade purchased from Sigma (Steinheim, Germany) or Merck (Darmstadt, Germany).

MS analyses were performed on an Orbitrap Fusion Lumos instrument (Thermo Fisher Scientific, Bremen, Germany) equipped with a robotic nanoflow ion source (TriVersa NanoMate, Advion

BioSciences, Ithaca, NY, USA) using chips with a spraying nozzle diameter of 5.5 μm. The back pressure was set at 1 psi. The ionization voltages were +1.3 kV and −1.9 kV in positive and negative modes, respectively, whereas it was +1.5 kV in acquisitions with polarity switching. The temperature of the ion transfer capillary was 260 °C. Acquisitions were performed at mass resolution Rm/z 200 = 240,000 in full scan mode. In the polarity switching method, spectra were acquired within the mass range of m/z 400–1300 from 0.2 to 0.6 min in the negative and from 0.8 to 1.2 min in the positive polarity mode. Phosphatidylethanolamine (PE), phosphatidylinositol (PI), phosphatidylserine (PS), phosphatidic acid (PA), phosphatidylglycerol (PG), cardiolipin (CL), and the lyso derivatives LPC, LPE, and LPI as well as ceramide (Cer), hexosyl ceramide (HexCer), and ceramide phosphoethanolamine (CerPE) were detected and quantified using the negative ion mode, whereas phosphatidylcholine (PC), diacylglycerol (DG), triacylglycerol (TG), and ergosteryl ester (EE) were detected and quantified using the positive ion mode.

For lipidomic measurements, 3 tumors per sample type were collected, immediately fresh-frozen in liquid nitrogen, and stored at −80 °C. For lipid extraction the samples were subjected to a one-phase methanolic lipid extraction. The tumors were sonicated, depending on the relative tumor size, in 150 – 300 μL methanol containing 0.001% butylated hydroxytoluene (as an antioxidant) in a bath sonicator for 5 min, then shaken for 5 min and centrifuged at 10,000 g for 5 min. The supernatant was transferred into a new Eppendorf tube and stored at −20 °C until MS analysis.

For MS measurements, 8-12 μL lipid extract was further diluted with 110 μL infusion solvent mixture (chloroform:methanol:iso-propanol 1:2:1, by vol.), which was spiked with an internal standard mix (Avanti Polar Lipids, Alabaster, AL, USA; **Supplementary Table S4**). Next, the mixture was halved, and 5% dimethylformamide (additive for the negative ion mode) or 3 mM ammonium chloride (additive for the positive ion mode) were added to the split sample halves. 10 μL solution was infused and data were acquired for 2 min.

Raw MS spectra were converted to platform-independent mzML files, and lipid species were identified by LipidXplorer software (Herzog et al., 2011). Identification was made by matching the m/z values of their monoisotopic peaks to the corresponding elemental composition constraints. The mass tolerance was set to 3 ppm. Data files generated by LipidXplorer queries were further processed by self-developed Microsoft Excel macros. Quantification was made by comparing integrated MS1 signal intensities with those of the internal standards.

Lipid species were annotated with sum formulas according to the shorthand notation for lipid structures^23^. For glycero(phospho)lipids, e.g., PE(34:1), the total numbers of carbons followed by double bonds for all chains are indicated. For sphingolipids, the sum formula like CerPE(34:1:2) specifies first the total number of carbons in the long chain base and FA moiety, then the sum of double bonds in the long chain base and the FA moiety, followed by the sum of hydroxyl groups in the long chain base and the FA moiety. The entire lipidomics dataset is available in **Supplementary Table S1**.

### Statistics

All image analyses were performed using the ImageJ image processing software and the resulting data were evaluated in GraphPad Prism 10 (using a free trial) except for the statistics of lipidomic data that was done in Microsoft Excel. Box plots were generated to visualize the data, and significance values are indicated on the statistical test charts as follows: *: p<0.05; **: p<0.01; ***: p<0.001; ****: p<0.0001. For simplicity in the case of visualization of lipidomic data, significant differences are marked with one asterisk only (that means p<0,05) and representation of p value was not detailed further.

#### Determination of tumor volume

To quantify tumor volumes Z-stack images were processed in ImageJ. A custom macro was applied that was built up the following steps: Images were denoised by Gaussian blur 3D function, then a mask were created for each image slice to identify tumor areas based on appropriate color intensity in the green channel by using the built-in,,Otsu” auto-thresholding model. Then, 3D objects were reconstructed from these masked images, representing tumor regions, and their volumes were measured by 3D objects counter function. By summing these volumes, the total tumor volume was obtained.

#### Quantification of lipid and immunofluorescent stainings

Images for each staining types were analyzed in ImageJ, with specific macros (see below). In parallel the area of GFP positive tumor tissues were also measured. Then the extracted values were aggregated and normalized to the total area of GFP+ tumor tissues by using a Python script. The processed values were then analyzed in GraphPad Prism. Within the same genotype, a paired t-test was performed if the data followed a normal distribution; otherwise, a Wilcoxon signed-rank test was used. For statistical analysis Student’s unpaired T-tests or Mann-Whitney tests were applied in case of datasets with normal or non-normal distribution respectively.

For MDH lipid staining analysis, a macro was used to compare the total area of lipid droplets in the GFP-positive tumor areas across the different genotypes. The,,Otsu” auto-thresholding model was applied for measuring GFP-positive areas, while the,,MaxEntropy” auto-thresholding model was used for masking the MDH channel. Then, the masked MDH channel was quantified in 2D using the “Analyze Particles” plugin.

For quantification of apoptotic areas, a similar approach to lipid staining analysis was applied. However, threshold was set by a custom script written in ImageJ, which calculated the masked area based on the characteristic size of the stained regions, each image mask was validated manually before quantification. The “Otsu” auto-threshold was still applied to mask the GFP-positive tumor areas. The masked areas were then quantified using the “Analyze Particles” plugin.

For P-H3 immunostaining, a custom macro was developed to compare the number of dividing cell nuclei in GFP-positive areas between control RS and ACC RNAi RS genotypes.,,Otsu” auto-thresholding method was again applied to GFP-positive areas, while the,,Moments” thresholding model was used for P-H3 staining. Two image slices (specifically, the third slice from both the top and bottom of each Z-stack image) were quantified using the “Analyze Particles” plugin, and the values from both slices were averaged for further calculations.

For immunostaining of autophagic structures, the evaluation was performed in ImageJ. GFP-positive tumor areas were masked using the,,Default” auto-threshold, while the Atg8a channel was masked with the,,Moments” auto-threshold model. Each Z-stack image consisted of seven slices (with 0,35 μm slice width) and measurements were taken from the middle three slices (3, 4, and 5) to ensure accurate quantification. The autophagic structures within tumor areas were analyzed on these three slices using the “Measure” command, and the values were then averaged to obtain a final measurement.

## Funding & Acknowledgements

This research was funded by the National Research, Development and Innovation Office of Hungary (OTKA FK_142508 to ST, OTKA ANN 139553 to GB), the Hungarian Academy of Sciences (BO/00400/23 to ST), and the Excellence Fund of Eötvös Loránd University (EKA_2022/045-P302-1 to ST). FOF was supported by a Norwegian Research Council Grant “Decoding tumor cell invasive switching” Project No.: 324447. We thank the Single Cell Omics Advanced Core Facility staff of the HCEMM for help with their resources and their support. HCEMM has received funding from the EU’s Horizon 2020 research and innovation program under grant agreement No. 739593 and KIM NKFIA 2022-2.1.1-NL-2022-00005.

We thank Sarolta Pálfia and Ivett Répássy and Benedek Tóth for their technical assistance.

## Author contributions

DK, FOF and ST designed and provided the resources for the research. DK, BSB NN, MP, GB performed the experiments and analyzed the data. MP and GB performed the lipidomics analysis. BSB wrote scripts and algorithms in Python and ImageJ. DK, FOF and ST wrote the manuscript.

