## Supplementary Figures S1-2 for "Fatty acid synthesis supports tumor progression through keeping TORC1 receptive for Insulin/PI3K signaling"

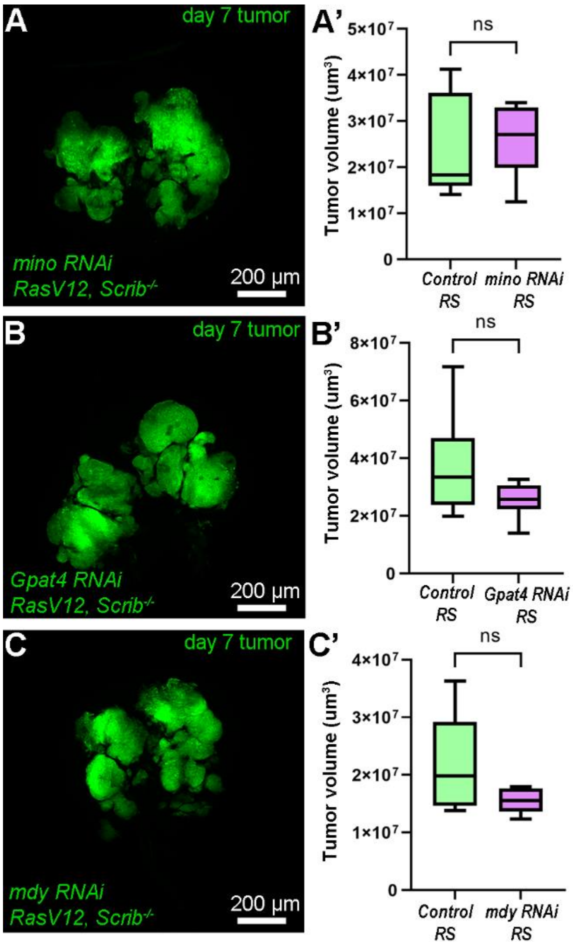


**Supplementary Fig. S1: Additional genetic screen data. (A-C)** Representative images at day 7 of mino **(A)**, Gpat4 **(B)** and mdy **(C)** deficient RS tumors from the RNAi screen. Quantification of respective tumor size data is shown on **A’-C’**. Please note that each RNAi dataset was compared to a dedicated control RS dataset that was collected and measured as part of the same experiment. 7-9 tumors were analyzed/genotypes, n=8 (control RS) and 9 mino RNAi RS) in A, 7 (control RS) and 8 (Gpat4 RS) in B, 7 (control RS) and 7 (mdy RNAi RS) in C. Mann-Whitney test (A), Student’s T-test (B, C), ns: non-significant.

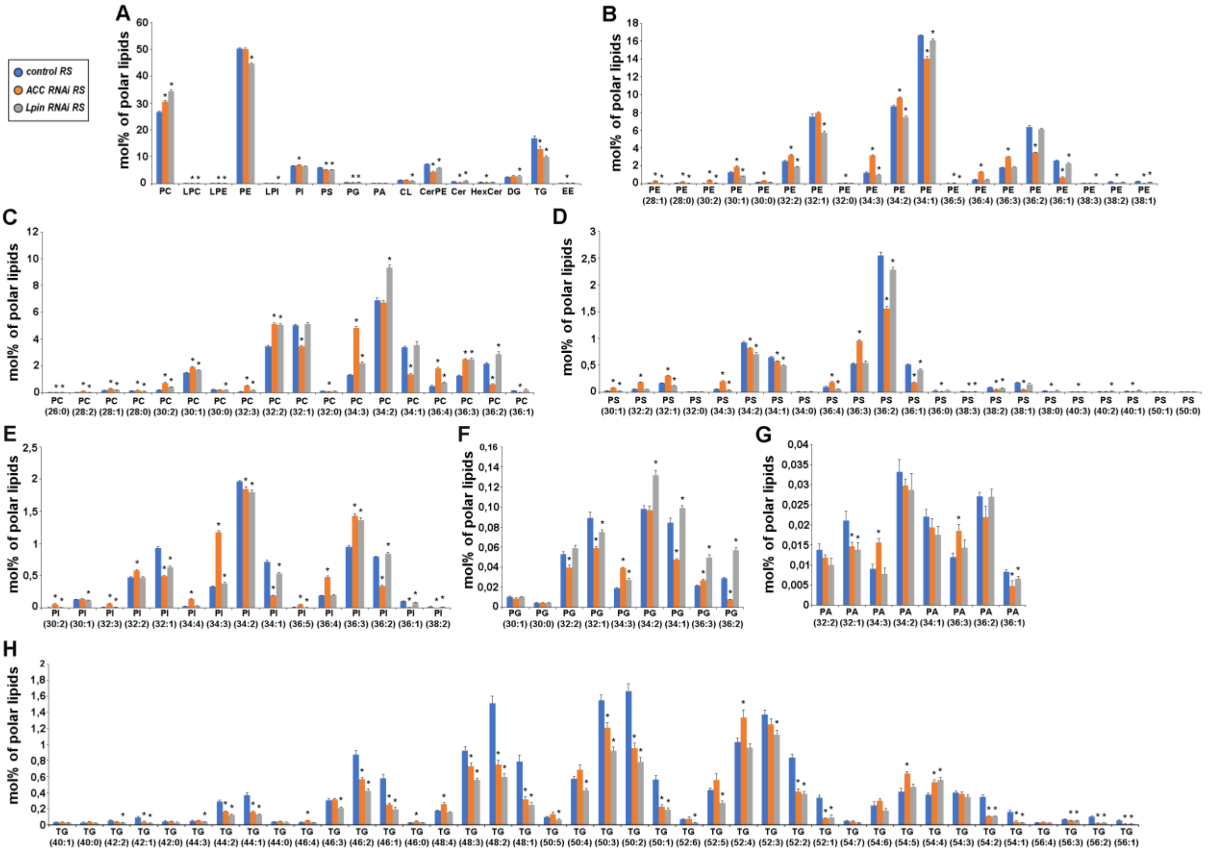


**Supplementary Fig. S2: Detailed lipid profiles of control and ACC or Lpin deficient RS tumors. (A)** Diagram representing the relative amounts of the major lipid classes. **(B-H)** Diagrams representing the relative amounts of all species of PE **(B)**, PC **(C)**, PS **(D)**, PI **(E)**, PG **(F)**; PA **(G)**; TG **(H)**. Statistical analysis on A-H was done by comparing values from ACC RNAi RS and Lpin RNAi RS to control RS data by using unpaired Student's T-test and asterisks (*) above the columns represent significant difference (p<0,05). 8-11 samples/genotypes were analyzed, n=10 (control RS), 8 (ACC RNAi RS),11 (Lpin RNAi RS). PC: Phosphatidylcholine; LPC: Lyso-phosphatidylcholine; LPE: Lyso-phosphatidylethanolamine; PE: Phosphatidylethanolamine; LPI: Lyso-Phosphatidylinositol; PI: Phosphatidylinositol; PS: Phosphatidylserine; PG; PA: Phosphatidic acid; CL: Cardiolipin; CerPE: Ceramide-phosphoethanolamine; Cer: Ceramide; HexCer: Hexosylceramide; DG: Diacylglycerol; TG: Triacylglycerol; EE: Cholesteryl-ester
