## Supplementary Table S4 for "Fatty acid synthesis supports tumor progression through keeping TORC1 receptive for Insulin/PI3K signaling"

| Lipid Class | Internal standard (IS) | Ion format | Conc of infusion (fmol/μL) |
|---|---|---|---|
| PC, LPC | PC(15:0/18:1-d7) | [M+H]+ | 480 |
| PE, LPE | PE(15:0/18:1-d7) | [M−H]- | 170 |
| PI, LPI | PI(15:0/18:1-d7) | [M−H]- | 119 |
| PS | PS(15:0/18:1-d7) | [M−H]- | 129 |
| PG | PG(15:0/18:1-d7) | [M−H]- | 26 |
| PA | PA(15:0/18:1-d7) | [M−H]- | 29 |
| CL | CL(tetra14:1) | $[M-2H]^{2-}$ | 16 |
| Cer | Cer(t18:1/16:0) | [M+Cl]− | 36 |
| HexCer | GluCer(d18:1/12:0) | [M+Cl]− | 31 |
| CerPE | CerPE(29:1:2) | [M−H]- | 340 |
| DG | DG(15:0/18:1-d7) | [M+NH4]+ | 34 |
| TG | TG(15:0/18:1-d7/15:0) | [M+NH4]+ | 124 |
| EE | Chold7E(16:1) | [M+NH4]+ | 153 |
